# Optimizing In Situ Proximity Ligation Assays for Mitochondria, ER, or MERC Markers in Skeletal Muscle Tissue and Cells

**DOI:** 10.1101/2023.05.20.541599

**Authors:** Dominique C. Stephens, Amber Crabtree, Heather K. Beasley, Edgar Garza-Lopez, Kit Neikirk, Margaret Mungai, Larry Vang, Zer Vue, Neng Vue, Andrea G. Marshall, Kyrin Turner, Jianqiang Shao, Sandra Murray, Jennifer A. Gaddy, Celestine Wanjalla, Jamaine Davis, Steven Damo, Antentor O. Hinton

**Author notes:** Corresponding Authors: Jamaine Davis, PhD, Meharry Medical College, Steven Damo, PhD, Fisk University, Antentor O. Hinton, Jr, PhD, Vanderbilt School of Medicine Basic Sciences. Co-first Authors (Dominique C. Stephens, Amber Crabtree, Heather K. Beasley).

## Abstract

Proximity ligation assays (PLA) use specific antibodies to detect endogenous protein-protein interactions. PLA is a highly useful biochemical technique that allows two proteins within close proximity to be visualized with fluorescent probes amplified by PCR. While this technique has gained prominence, the use of PLA in mouse skeletal muscle (SkM) is novel. In this article, we discuss how the PLA method can be used in SkM to study the protein-protein interactions within mitochondria-endoplasmic reticulum contact sites (MERCs).

**Tweetable Abstract:** Proximity Ligation Assays can be used in skeletal muscle tissue and myoblasts to explore the protein-protein interactions involved in MERC sites.

**Highlights:**

- Skeletal muscle tissue and cells are plated on glass coverslips for evaluation by proximity ligation assay (PLA).
- Following fixation, cells are probed and stained for Mfn1, Mfn2, mitochondria, and ER and imaged using fluorescence confocal microscopy.
- This method shows that PLA can be used in mouse SkM and is adaptable to other models.
- Protocol for detection of protein-protein interactions using PLA.

**Graphical Abstract:** 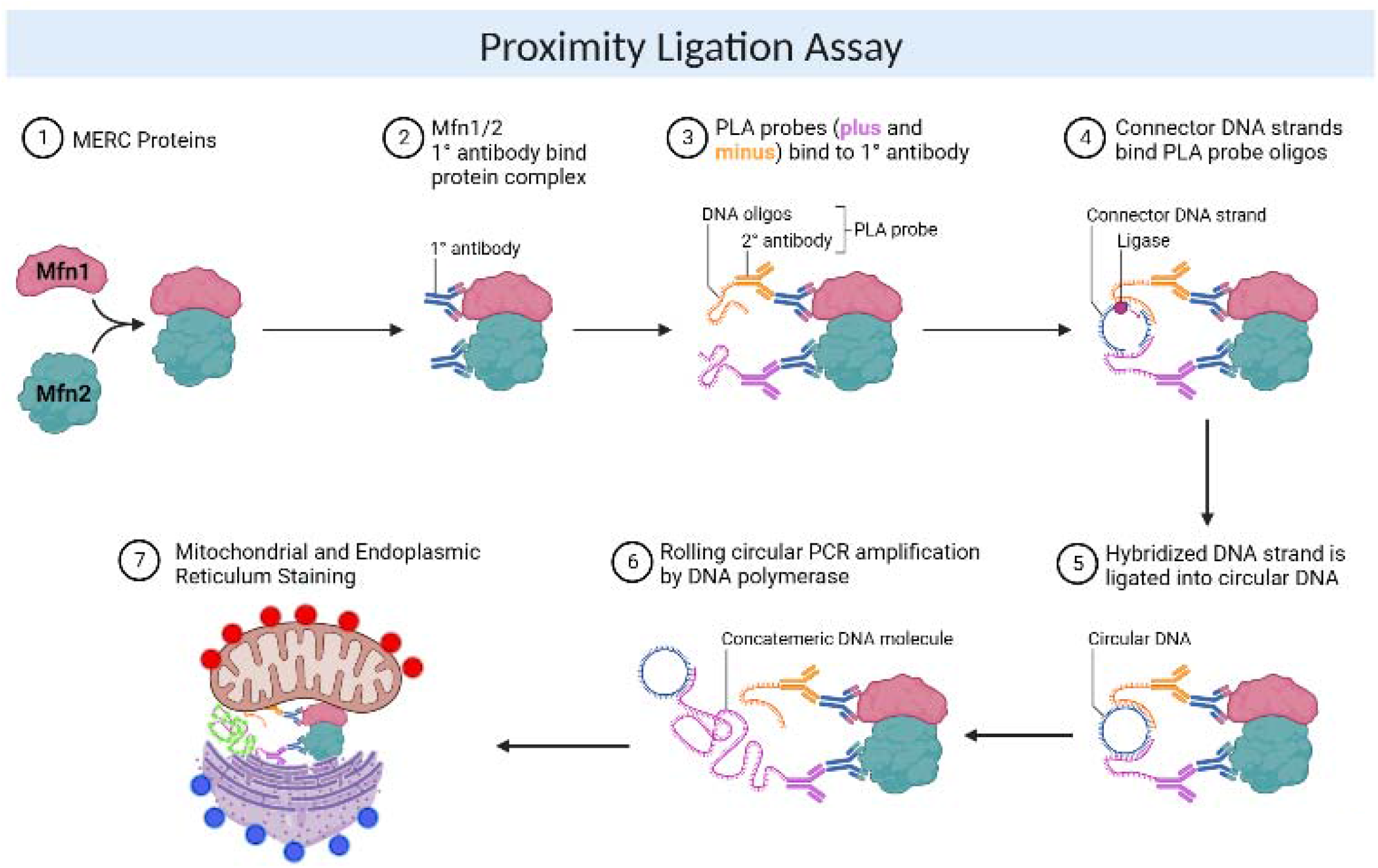

| **Key Resources Table** |  |  |
| --- | --- | --- |
| **<u>Reagent or Resource</u>** | **<u>Source</u>** | **<u>Identifier</u>** |
| **Antibodies** |  |  |
| MFN1 Monoclonal antibody | abcam | ab126575 |
| MFN2 Monoclonal antibody | abcam | ab124773 |
| **Stain** |  |  |
| MitoStain | Bio-Rad | 1351321 |
| Cytopainter | abcam | ab139482 |
| **Chemicals** |  |  |
| Triton X-100 | Thermo Scientific | 9002-93-1 |
| Paraformaldehyde | Thermo Scientific | 50-00-0 |
| PBS, pH 7.4 | Thermo Scientific | 10010023 |
| Duolink® In Situ Wash Buffers | Sigma-Aldrich | DUO82049 |
| NaCl (5 M) (If not purchasing Duolink® In Situ Wash Buffers ) | Thermo Scientific | AM9760G |
| TRIS, 1.0M., pH 7.4 (If not purchasing Duolink® In Situ Wash Buffers) | Thermo Scientific | J60202-K2 |
| UltraPure™ 1 M Tris-HCl Buffer, pH 7.5 (If not purchasing Duolink® In Situ Wash Buffers) | Thermo Scientific | 15567027 |
| TWEEN® 20 0 (If not purchasing Duolink® In Situ Wash Buffers) | Sigma-Aldrich | P1379 |
| **Quantification and Software** |  |  |
| GraphPad | GraphPad Software, San Diego, California USA | www.graphpad.com |
| Image J | Schindelin et al.5 | https://imagej.net/ |
| **Critical Commercial Assays** |  |  |
| Duolink® PLA Kit | Sigma-Aldrich | DUO92008 |
| Duolink® In Situ Red Starter Kit Mouse/Rabbit | Sigma-Aldrich | DUO92101 |
| **Materials** |  |  |
| Humidity Chamber | Biogear | BGHC-024 |
| Leica Microm Cryostat I HM505E | Leica | 8243-30-1000 |
| DWK Life Sciences Wheaton™ Staining Jar | Fisherscientific | 22-312143 |

## Introduction

Protein-protein interactions are essential for cellular communication, as it is the primary means of communication between organs, tissues, cells, and organelles ^1^. Various methods are often used to o study these interactions including co-immunoprecipitation (co-IP), pull-down assays, crosslinking, label transfer, and far western blotting. However, Proximity Ligation Assays (PLA) have emerged as an enhanced method for detecting protein-protein interactions. PLA is highly specific and detects pairs of oligonucleotide-conjugated antibodies to identify interactions utilizing fluorescence microscopy ^1^.

PLA takes advantage of close proximity binding (with distances up to 40 nm apart). Primary antibodies of two different species are used to target the endogenous proteins of interest ^2^. A pair of oligonucleotide-labeled secondary antibodies will then generate a signal if they are within close proximity of each other ^2^. These highly specific antibodies can bind to different epitopes of the same protein or to two proteins in a complex and can provide localized detection, visualization, and quantification of the protein(s) of interest ^1^. There are many application possibilities with PLA, including adherent cell lines, cytospin preparations, and tissues, including frozen or paraffin-embedded patient samples ^3^. However, it should be noted that cells and tissues for this assay must be fixed and permeabilized prior to the use of PLA antibodies ^3^.

Another advanced method often used that is similar to PLA is FRET (Fluorescence Resonance Energy Transfer) which analyzes interactions between two fluorescently tagged proteins ^4^. When comparing the two, FRET is a cheaper option that can be as equally informative as PLA. This is due to the costlier price tag of PLA probes in comparison to FRET fluorophore-tagged secondary antibodies. Additionally, although FRET and PLA are both antibody-based methods, FRET indicates protein interactions within 10 nm, whereas PLA can be used for protein interactions that are up to 40 nm apart ^5^. This increased distance allows for PLA to detect proteins that are in close proximity to each other but might not be physically interacting with each other at that time. Perhaps the main distinguishing factor between PLA and FRET is the ability to quantify the signal amplification in PLA whereas FRET can only be used as a qualitative measure. Additionally, PLA is not limited to a low signal-to-noise ratio associated with imaging that is associated with FRET ^4^. This has caused PLA to arise as a fundamental alternative to FRET that can be important in elucidating the spatiotemporal dynamics and interactions of proteins.

In recent years, PLA has been used to explore mitochondria-endoplasmic reticulum contact sites (MERCs) to fully understand protein-protein interactions between organelles in contact with each other ^1^. Two important mitochondrial proteins are Mitofusins 1 and 2 (Mfn1 and Mfn2), which are both located on the outer mitochondrial membrane (OMM) but have different functions ^6^. Mfn1 mainly functions to facilitate organelle docking and fusion of the OMM. Mfn2, on the other hand, is highly expressed at MERCs where it controls organelle tethering ^7^. The distinct functions of Mfn1 and Mfn2 as well as their colocalization make them ideal to study MERC protein-protein interactions in skeletal muscles using PLA. However, it should be noted that PLA is only as accurate and effective as the antibodies and markers used. With the selection of adequate antibodies and markers for PLA, this protocol can be used in a variety of ways. Here, we describe the methodology of PLA for staining mouse SkM mitochondria and ER through probing Mfn1 and Mfn2.

## Materials and Methods

Here, we describe how to use a Duolink^®^ PLA fluorescence protocol in primary myoblasts ^8^ to probe for Mfn1 and Mfn2. Due to long incubation times, this procedure typically takes up to 1-2 days, but once completed, imaging can proceed shortly thereafter depending on the mounting media and typically samples should ideally be imaged within a few days ^9^. Notably, this procedure can also be utilized for skeletal muscle tissue in addition to myoblasts.

Animal work followed the University of Iowa Animal Care and Use Committee guidelines and had approval. Animals were housed at 22°C with a 12 h light, 12 h dark cycle with free access to water and standard chow. The generation of 9-month *Opa1*^*fl/fl*^ and OPA1 smKO (*OPA1*^*flox/flox*^ HSA-CreER^T2^) C57Bl/6J mice was performed per prior established protocols ^10^. Briefly, the null Opa1 allele was generated by crossing Flox-neo mice with EIIa-Cre transgenic mice, resulting in a stop codon after exon 9 due to a frameshift ^11^. Once the gastrocnemius muscle was excised, the protocol below was slightly modified but utilized as described.

**Note:** This protocol is a modified version of the manufacturer’s protocol. The manufacturer’s protocol can be found at: https://www.sigmaaldrich.com/US/en/technical-documents/protocol/protein-biology/protein-and-nucleic-acid-interactions/duolink-fluorescence-user-manual

### Before you Begin

#### Solutions Needed

Please note that most of the solutions listed are contained within the Duolink® PLA Kit (Sigma DUO92008). Upon receiving the PLA kit (Sigma DUO92008), note that the components have 3 different storage conditions. Be careful to store each component as recommended. Any indicated volumes are for 40 μL of solution to properly cover a 1 cm^2^ sample on a slide. Adjust the volumes accordingly to the number of samples and area of coverage used.

#### Fixation solution

| Reagent | Stock Concentration | Final Concentration | Amount |
| --- | --- | --- | --- |
| Paraformaldehyde | 16% | 4% | 10mL |
| PBS | 2X | 1X | 20mL |
| dd H<sub>2</sub>O | N/A | N/A | 10mL |
| Total | N/A | N/A | 40mL |

**CRITICAL**: The purity of water can affect the signal of PLA, so high-purity water (dd H_2_O) should consistently be used across this protocol.

#### Wash solution

| Reagent | Amount Needed |
| --- | --- |
| PBS (Dulbecco's pH 7.4) | 500mL |

**Note:** You will need to verify the proper pH is attained.

#### Permeability Solution (0.2%)

| Reagent | Amount Added |
| --- | --- |
| Triton X-100 (100%) | 20.1uL |
| PBS | 10mL |
| Total | 10.02 mL |

The Antibody Diluent will come in the Duolink® In Situ PLA® Probe kit ready to use.

**Note:** the diluent should be stored at 4 °C upon arrival. Vortex the diluent before use.

The <u>Blocking Solution</u> will also be supplied in the Duolink® In Situ PLA® Probe kit, ready to use, and should be stored at 4 °C upon arrival. Vortex before use.The Antibody Diluent will come in the Duolink® In Situ PLA® Probe kit ready to use.

#### Mfn1 and Mfn2 Primary Antibodies

Antibodies should be diluted in Duolink® Antibody Diluent to make working solutions.

**Note:** Antibodies should only be used that have been tested for optimal conditions (antigen retrieval method; permeabilization conditions, if required; antibody concentration).

#### PLA Probes (1:5 Dilution)

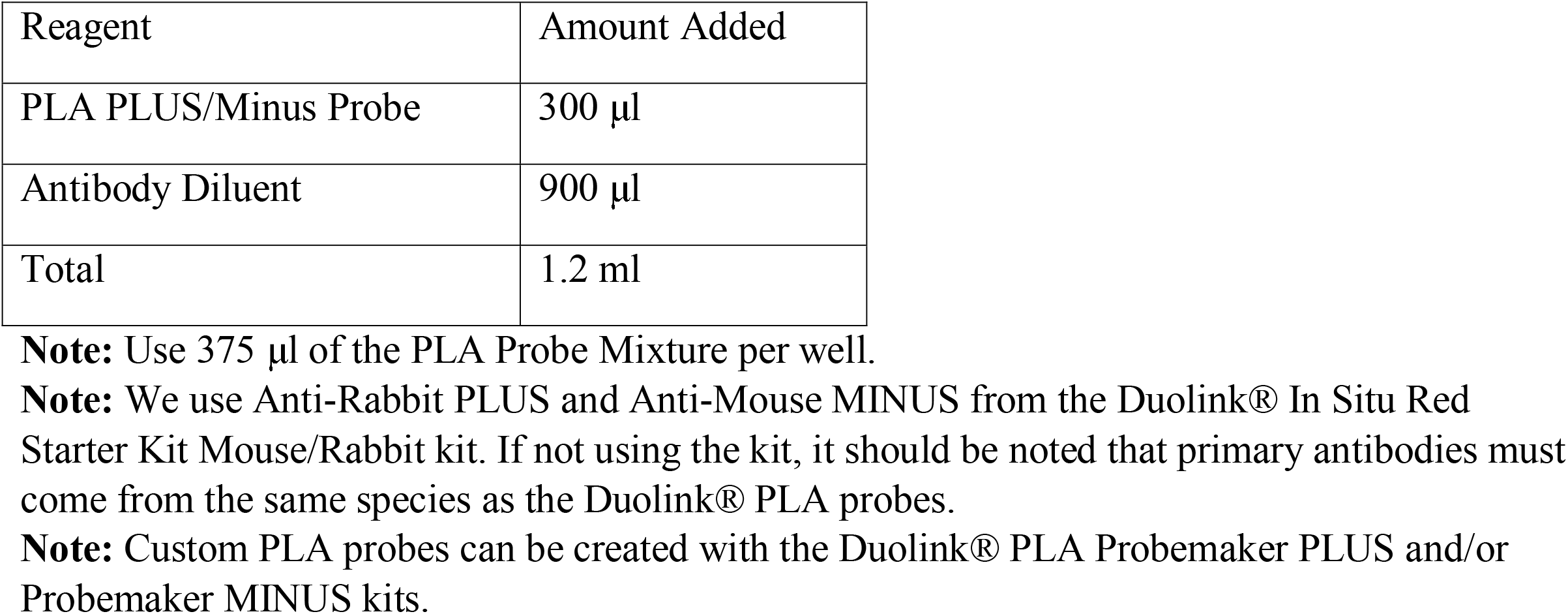

#### WASH Buffer

| Reagent | Amount Added |
| --- | --- |
| Washer Buffer A/B | 1 Pouch |
| dd H<sub>2</sub>O | 1 L |
| Total | 1 L |

**Note:** WASH Buffers, once resuspended, should be stored at 4 °C upon arrival.

**N**OTE: Wash Buffers may also be manually made through the two tables below, if not commercially bought. If prepared, filter with a 0.45 μm filter and store at 4 °C.

#### WASH Buffer A

| Reagent | Amount Added |
| --- | --- |
| Tween 20 | 0.5 mL |
| NaCl (5 M) | 30 mL |
| Tris (1 M), pH 7.4 | 10 mL |
| dd H<sub>2</sub>O | 959.5 mL |
| Total | 1 L |

#### WASH Buffer B

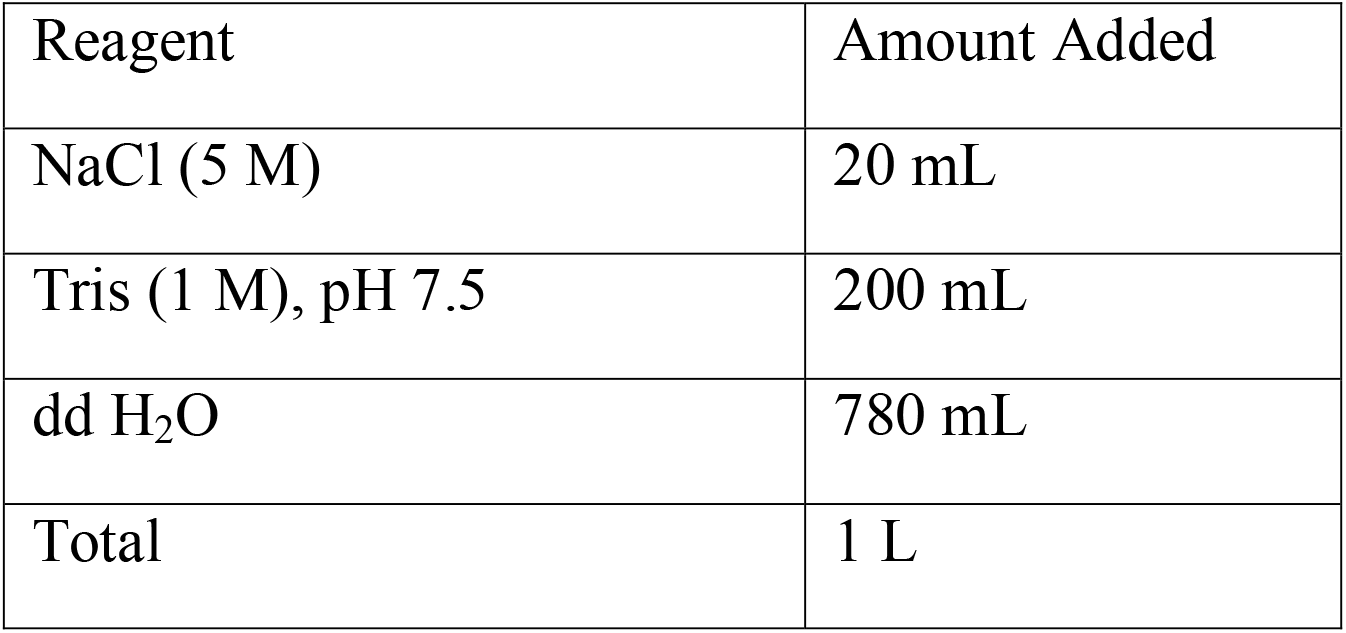

#### 1X Ligation Buffer (1:5 Dilution)

| Reagent | Amount Added |
| --- | --- |
| Ligation 5x Buffer | 8 µl |
| dd H<sub>2</sub>O | 32 µl |
| Total | 40 µl |

#### Working Ligase Buffer

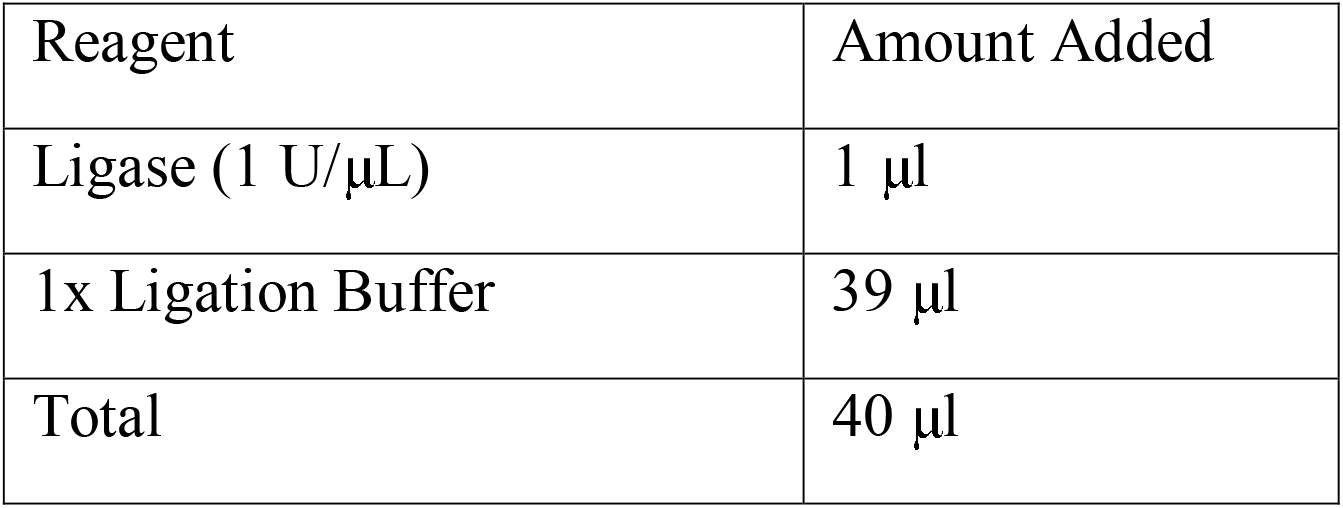

**CRITICAL:** Ligation buffer contains Dithiothreitol (DTT). No DTT precipitates should be visible prior to the preparation of the ligase reaction buffer to ensure

#### Amplification Buffer (1:5 Dilution)

| Reagent | Amount Added |
| --- | --- |
| 5x Amplification Buffer | 8 μl |
| dd H<sub>2</sub>O | 32 μl |
| Total | 40 μl |

#### Working Amplification Solution (1:80 Dilution)

| Reagent | Amount Added |
| --- | --- |
| Amplification Buffer | 39.5 μl |
| Polymerase | 0.5 μl |
| Total | 40 μl |

#### 0.01x Wash Buffer B

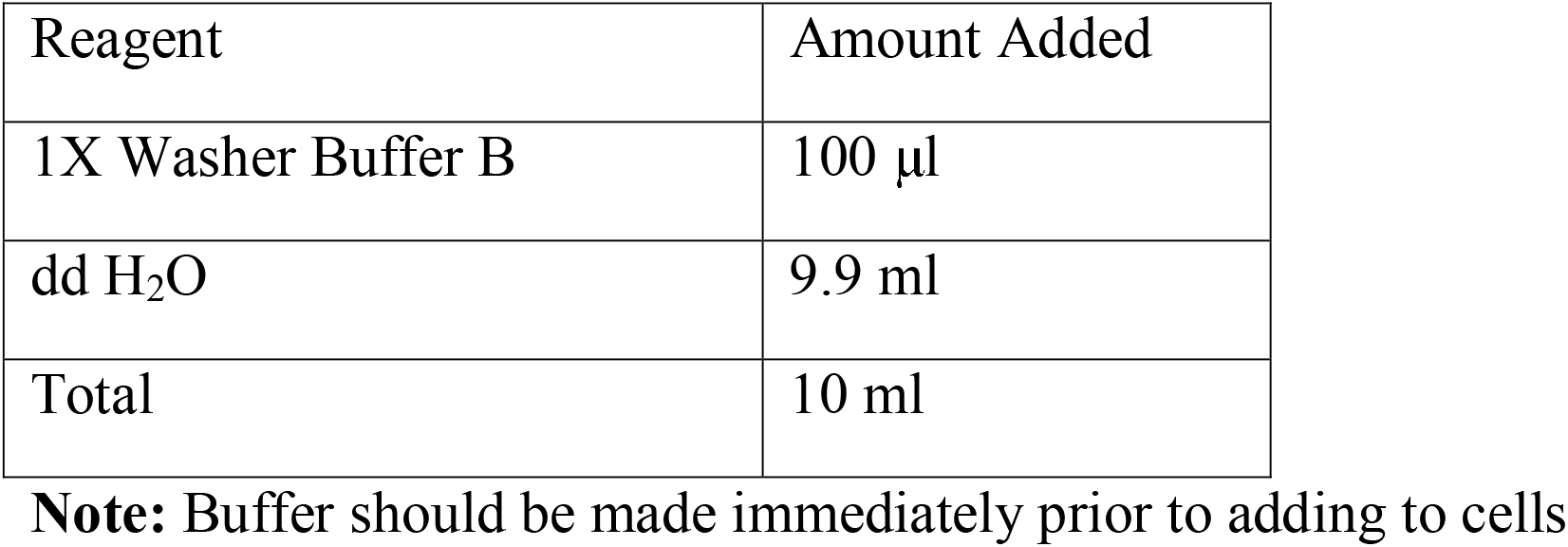

#### Duolink® Mounting Media

will also come ready to use with the Duolink® In Situ PLA® Probe kit and should be stored at 4 °C upon arrival.

**Note**: Three technical controls should be included in every Duolink® PLA experiment. These include: the omission of each primary antibody separately (used to detect non-specific binding of each primary antibody; helpful in determining optimal primary antibody titer) and the omission of all primary antibodies (used to detect non-specific binding of the Duolink® PLA Probes in the system).

### Preparation of Tissues and Cells (Day 0)

This is a highly adaptable protocol that can use tissues or cells prepared through several mechanisms. The preparation we have used for cells and tissues is offered below:

1. Day 0: Cells
  a. If using cells, isolate myoblasts and differentiate into myotubes in accordance with prior protocols^8^ beginning 2 weeks prior to this protocol.
  b. Plate 150,000 cells per borosilicate chamber slide at 70% confluency. **CRITICAL**: Avoid overconfluency which can lead to false positives in the PLA process.
  c. Pipette 500 μl media, enough to cover cells, in the chamber slides.
2. Day 0: Cryo-Sections Tissue
  a. Following IACUC guidelines, euthanize mice.
  b. Collect muscle tissue from the gastrocnemius muscles in both legs.
  c. Incubate in fixation solution (500 μl per chamber) for 24 hours at 37°C.
  d. Wash 3 times with PBS.
  e. Place the tissue sample in an optimal cutting temperature (OCT) medium following fixation.
  f. Place tissue samples on a cryostat for approximately 2-3 minutes until freezing for sectioning
  g. Upon freezing, perform slicing on the cryostat
  h. Place tissues sections on slides
3. Day 0: Paraffin-Sections Tissue
  a. Following paraffin-embedded samples, sections were dried 12-16 hours (overnight) on a 45°C slide warmer.
  b. Prior to decalcification, the slides were warmed to the melting point in a 60°C oven, which will depend on the number of slides being utilized.
  c. The warm slides were deparaffinized in a stain jar filled with xylenes for incubation 3 times (5 min each). Another stain jar was filled with 100% ethanol and slides were incubated twice (1 min), then in another stain jar with 95% ethanol twice (1 min), and finally a stain jar with distilled water.

**CRITICAL**: All incubations should be performed in an incubator or humid environment and samples should be hydrated enough so that they do not dry out during this process.

#### Experimental Protocol

1. Day 1: **Timing: 2 hrs**. Fixing Cells **NOTE**: If using tissue, omit A-D, but all other steps are the same unless otherwise noted.
  a. Aspirate media from the cells **NOTE**: Pipettes or low speed aspirators can decant this solution.
  b. Incubate in fixation solution (500 μl per chamber) for 10 mins at 37°C.
  c. Remove the fixation solution.
  d. Wash 3 times with PBS. Permeabilize cells/slides
  e. Incubate cells or slides in permeabilization solution (500 μl per chamber) for 10 mins.
  f. Wash the cells 3 times with PBS. **NOTE**: Rocking during wash steps are not necessary in our experience.

#### Duolink® PLA Protocol

Prior to beginning this protocol, samples should be processed as mentioned above.

<u>Blocking</u>

g. Vortex the Duolink® Blocking Solution prior to adding to the sample.
h. Cover the sample with 40 μL Duolink® Blocking Solution.
i. Incubate for 1 hour at 37 °C in a humid chamber. **CRITICAL:** Do not let the sample/slide dry prior to adding the antibody.
j. Vortex the Duolink® Antibody Diluent.
k. Dilute your primary antibody (Mfn1/mfn2 antibodies) to the necessary concentration.
l. Aspirate the Duolink® Blocking Solution from the slides.
m. Add the primary antibody solution to each sample and incubate at 4°C. **Note:** Slides should be incubated using the appropriate temperature and time for the primary antibody that produces optimal results.

**CRITICAL**: IgG antibody pairs must also be from two different species (mouse, rabbit, goat or human)

2. Day 2:

**Timing: 6 hrs**.

<u>Duolink® PLA Probe Incubation</u>

a. Vortex both PLA probes (PLUS and MINUS).
b. Dilute the probes (PLUS and MINUS) 1:5 with Duolink® Antibody Diluent
c. Aspirate the primary antibody solution from the slides.
d. Wash the slides for 5 minutes at room temperature in 1X Wash Buffer A 2x.
e. Aspirate the excess wash buffer.

**CRITICAL**: Ensure minimal wash buffer remains on the sample, as it may dilute future solutions.

f. Apply The PLA Probe Solution.
g. Incubate for 1 hour at 37 °C in humid chamber.

<u>Ligation</u>

**Note:** Do not add the Ligase to the ligation buffer until immediately before adding it to the sample.

h. Dilute the 5x Duolink® Ligation buffer 1:5.
  i. Use sterile filtered or Milli-Q® water.
  ii. Mix the solution thoroughly.

**Note:** Prepare enough solutions needed for all samples.

i. Aspirate the PLA probe solution from the slides.
j. Wash the slides/cells with enough 1X Wash Buffer A to cover the slide (∼1 mL). Wash 2 times at room temperature for 5 mins each.
k. Remove the Ligase from the freezer.

**Note:** Use a freezer block for the ligase tube.

l. Dilute the Ligase 1:40.
  iii. Use the 1X Ligation Buffer from the last step.
m. Aspirate the wash buffer.
n. Add the ligation solution to the sample.
o. Incubate at 37 °C for 30 mins in humid chamber.

**TIME CRITICAL STEP:** Ensure ligation is precise in timing, as too long can result in false positive PLA signal.

<u>Amplification</u>

**Note:** Do not add the Polymerase until immediately before adding it to the sample. Remember to protect the amplification buffer from light.

p. Dilute the 5X amplification buffer 1:5.
  iv. Use sterile filtered or Milli-Q® water

**Note:** Make sure to make enough diluted amplification buffer to cover all samples.

q. Aspirate the ligation solution from the slides.
r. Wash the slides in 1X WASH Buffer A for 5 mins at room temperature two times.
s. Remove the Polymerase from the freezer using a freezer block.
t. Dilute the Polymerase 1:80.
  v. Use the 1X Amplification Buffer from the last step for dilution.
u. Mix thoroughly.
v. Aspirate the excess WASH buffer.
w. Add the amplification solution to the sample.
x. Incubate at 37 °C for 100 mins in humid chamber.

**TIME CRITICAL STEP:** Ensure 100 minutes for amplification are performed.

**CRITICAL**: Ensure this amplification step is protected from light.

**Note:** Always protect the slides from light.

y. Aspirate the mounting media or PBS from the slides.
z. Wash the slides with Wash Buffer B at 20 °C to 25 °C (room temperature) for 10 mins two times.
aa. Wash slides for 60 seconds.
  a. Use 0.01X Wash Buffer B

<u>Mitochondria and Endoplasmic Reticulum Staining</u>

Mfn2 is an important MERC protein in addition to being a mitochondrial protein, while Mfn1 is exclusively a mitochondrial fusion protein. Staining of both the mitochondria and ER is necessary to understand the protein-protein interactions within MERC regions to co-localize with puncta and show organelles. While this step is not necessary, it can aid in visualization.

bb. Pre-warm the working stain solution [MitoStain and ER-Tracker].
cc. Aspirate any residual solution from the samples.
dd. Add the working stain solution.
ee. Incubate the samples for 30 mins at 37°C in a humid chamber.
ff. Aspirate the stain and replace it with warm mounting media.

<u>Final Washes</u>

gg. Proceed to imaging preparation.

**CRITICAL**: Coverslip edges are prone to drying, so avoid coverslips or tissue samples should not be dried until following the final wash buffer prior to mounting ^9^.

<u>Preparation for Imaging</u>

**Note:** Duolink® In Situ Mounting Media with DAPI will not solidify. The cover slip can be sealed using clear nail polish around the edges.

hh. Aspirate the wash buffer from the previous step from the slides.
ii. Add the smallest amount necessary to mount Duolink® In Situ Mounting Medium with DAPI
jj. Mount the slides with a coverslip.
kk. Incubate mounted slides for 15 mins
ll. Perform analysis *via* a confocal or fluorescence microscope. Here, we use an excitation of 594nm and an emission of 624nm.

**Note:** Following imaging, the slides can be stored for up to 6 months at -20 °C and up to 4 days at 4 °C. Make sure that the slides are protected from light.

### Expected Outcomes

As an example, we sought to understand how optic atrophy protein 1 (OPA1)-deficiency causes alteration in Mfn1-Mfn2 interactions. OPA1 is a common mitochondrial fusion protein that is understood to interact with Mfn2 to modulate MERC spacing ^12^. Here, we performed a knockdown of OPA1 through satellite cell isolation of *Opa1*^*fl/fl*^ double-floxed mice. BD Matrigel-coated dishes and differentiated into myoblasts and adenovirus expressing GFP-Cre were utilized to cause OPA1 deficiency as per established protocols ^10^. First, we used total internal reflection fluorescence (TIRF) to look at Mfn1-Mfn2 interactions. In myoblasts, interactions were visualized in both WT myoblasts and *Opa1*-deficient myoblasts (Figure 1A-F). Quantification further validated that the loss of *Opa1* caused increased Mfn1-Mfn2 interactions (Figure 1G). From there, we looked at confocal PLA. Nuclei were also labeled with 4′,6-diamidino-2-phenylindole (Figure 2A-B), and Mfn1-Mfn2 interactions were tagged (Figure 2C-D). Merging channels allowed the understanding of Mfn1-Mfn2 interactions relative to the nucleus following the loss of *Opa1* (Figure 2E-F). Quantification showed that there was a significant increase in Mfn1 and Mfn2 interaction puncta following the loss of *Opa1* (Figure 2G).

**Figure 1:**
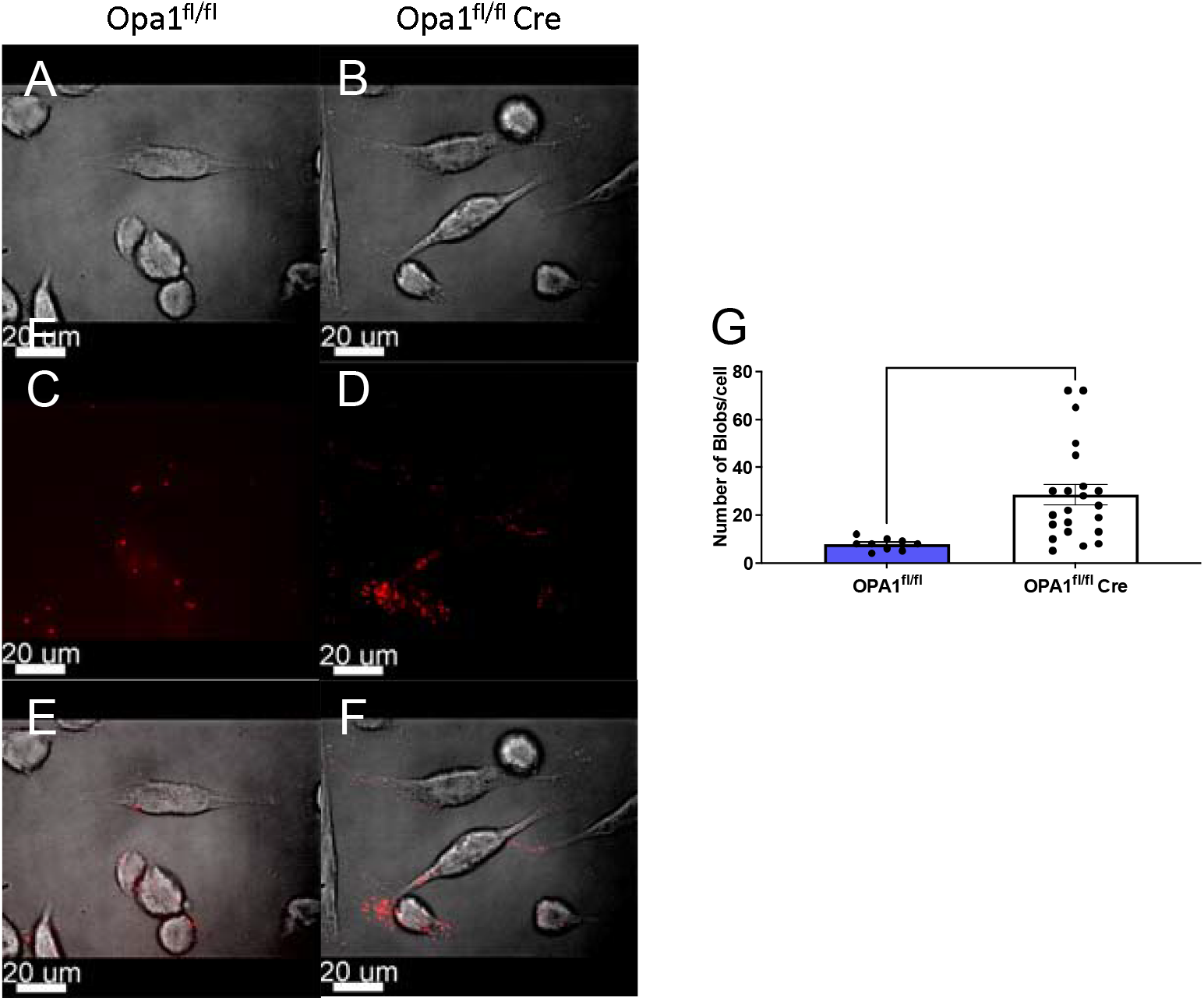
Total internal reflection fluorescence (TIRF) microscopy showing Mfn1–Mfn2 interactions (red punctae) in murine myoblasts. (A) Isolated myoblasts in Opa1 and (B) Opa1^fl/fl^Cre. (C) Isolated PLA fluorescence in Opa1 and (D) Opa1^fl/fl^Cre. (E) TIRF myoblasts overlaid with proximity ligation assay (PLA) fluorescence in Opa1 and (F) Opa1^fl/fl^Cre. (G) Quantification of normalized PLA Mfn1-Mfn2 interactions in Opa1 and Opa1^fl/fl^Cre through TIRF. Dots represent the sample number, which was done in triplicates. An unpaired t-test was used to measure the statistical difference and** indicates p<0.01.

**Figure 2:**
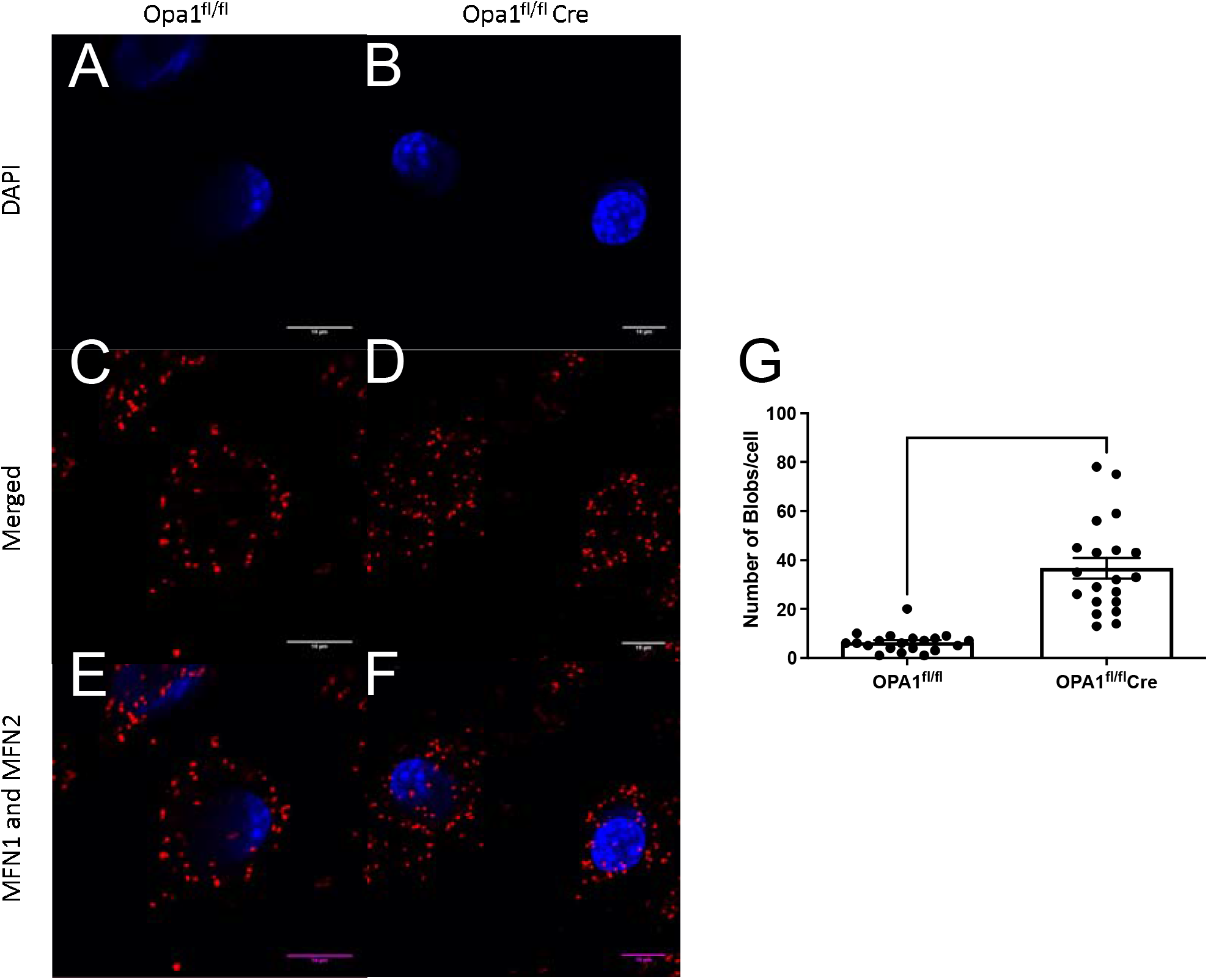
*In situ* confocal proximity ligation assay (PLA) highlighting Mfn1–Mfn2 interactions (red punctae) in murine myoblasts. (A) Nuclei are labeled with 4′,6-diamidino-2-phenylindole (DAPI, blue) in Opa1^fl/fl^ and (B) Opa1^fl/fl^Cre. (C) Mfn1-and Mfn2 interactions in Opa1^fl/fl^ and (D) Opa1^fl/fl^Cre. relative to nuclei. (E) Merged channel showing PLA Mfn1-Mfn2 interactions in Opa1 and (F) Opa1^fl/fl^Cre. (G) Quantification of normalized PLA Mfn1-Mfn2 interactions in Opa1 and Opa1^fl/fl^Cre. Dots represent the sample number, which was done in triplicates. An unpaired t-test was used to measure the statistical difference and **** indicates p<0.0001.

Here we used this same protocol for a tissue sample (Figure 3A-B). We found that Mfn1-Mfn2 localization could also be observed (Figure 3C-D). We then analyzed isolated PLA puncta of Mfn1-Mfn2 interactions (Figure 3E-F). In SkM, similar to myoblasts, we were able to verify that Opa1 loss causes an increase in Mfn1-Mfn2 interactions (Figure 3G).

**Figure 3:**
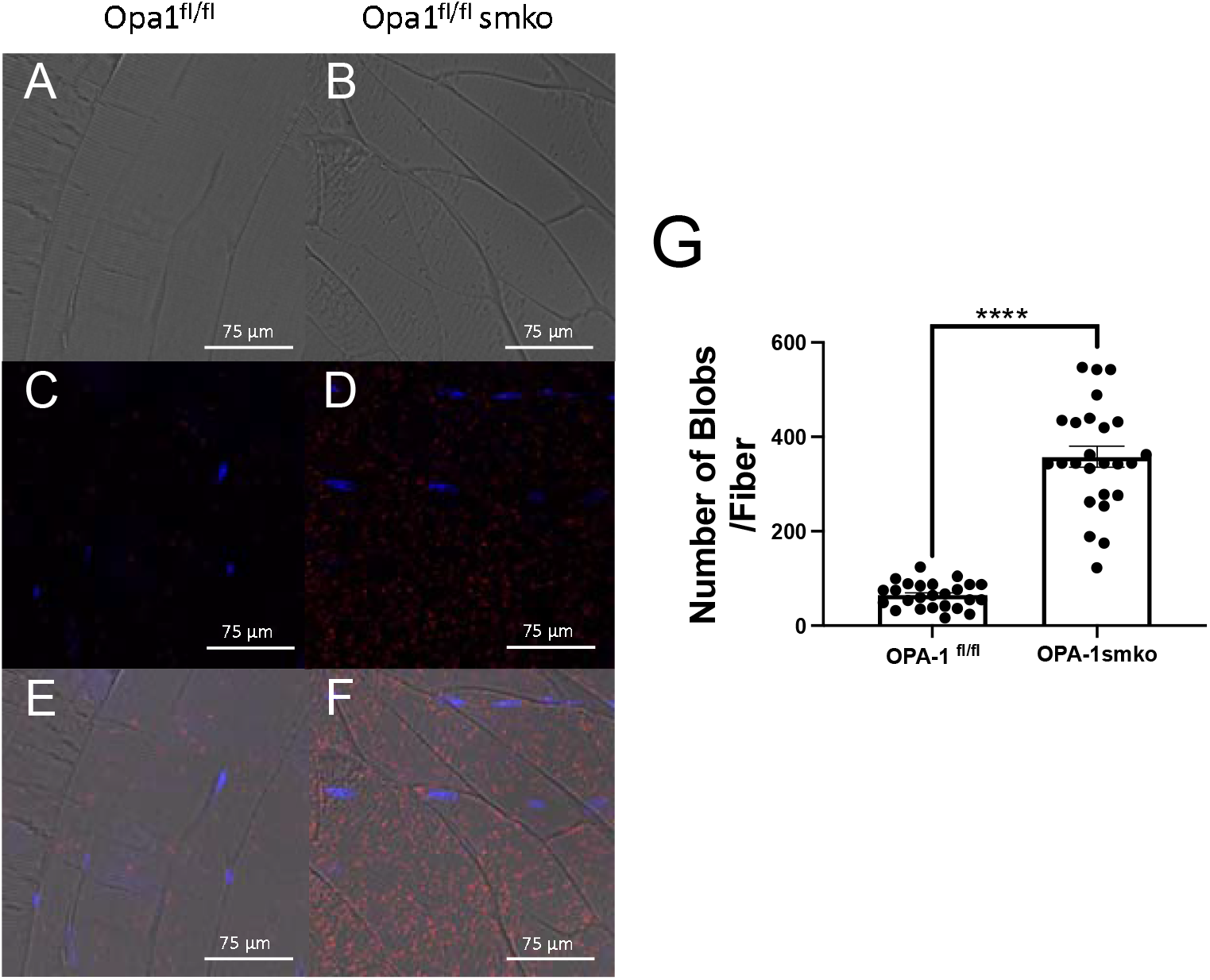
*In situ* PLA confocal visualization of murine gastrocnemius skeletal muscle in 40-week-old WT and *OPA1* smKO mice. Mfn1–Mfn2 interactions (red punctae) are shown and nuclei are labeled with 4′,6-diamidino-2-phenylindole (DAPI, blue). (A) Representative image of murine skeletal muscle in Opa1^fl/fl^ (B) and Opa1^fl/fl^ smKO. (C) Isolated PLA fluorescence in Opa1 and (D) Opa1^fl/fl^smKO. (E) Fluorescence in Opa1^fl/fl^ and (F) Opa1^fl/fl^smKO showing Mfn1-and Mfn2 interactions relative to nuclei overlaid on skeletal muscle. (G) Quantification of normalized PLA Mfn1-Mfn2 interactions in Opa1 and Opa1^fl/fl^smKO. Dots represent the sample number, which was done in triplicates. An unpaired t-test was used to measure the statistical difference and **** indicates p<0.0001.

### Quantification and Statistical Analysis

After imaging, blobs of PLA were quantified relative to the cell or fiber to provide standardization. Quantification can be done through ImageJ with auto-threshold and consistent settings between groups, as per established protocols ^9^. With consistent acquisition parameters between samples, a multitude of alternative tools can be used, such as Blobfinder (http://www.cb.uu.se/~amin/BlobFinder/). GraphPad Prism version 8.4.0 (GraphPad Software, La Jolla, CA) was used to perform students’ T-tests to measure statistical significance.

### Limitations

One of the largest limitations is that false positives may occur with PLA, due to cross-reactivity. Specifically, several steps should be taken regarding non-specific binding or false positives. Importantly, for tissues, one slide should serve as a negative devoid of both primary antibodies and inactivated antibodies, to detect any non-specific binding of the Duolink PLA probes. Recently, using Flag-tags and PLA together, past studies have stained for ER and mitochondria prior to performing PLA to verify through confocal microscopy the localization of proteins in specific regions ^13^. Staining for ER and mitochondria can therefore provide validation that false positives to regions outside of interest or in MERCs zones are not happening. Multicolor 3D imaging pipelines have also been established for the distribution of proteins through PLA in bone marrow ^14^. However, spatial changes in protein distribution being concomitant with changes in morphology are inhibited by the lack of protocols to perform both EM and PLA on the same sample.

Similarly positive controls are also important and can be time-consuming. Given the specificity of PLA on precise timing, steps need to be taken to ensure lack of signal arises due to experimental conditions and not improper technique, which can be difficult to discern. Positive controls can be differentiated including validated interactions can be important to validate that probes are effective. Together, these positive and negative controls while not providing certainty, can aid in the confirmation of mitochondria and ER morphology and localization for Mfn1 and Mfn2, or other mitochondrial markers.

### Trouble Shooting

Here we provide some common issues. This list is non-comprehensive and a useful resource is offered by Sigma Aldrich (https://www.sigmaaldrich.com/GB/en/technical-documents/technical-article/protein-biology/protein-and-nucleic-acid-interactions/duolink-troubleshooting-guide, accessed 5/1/2023).

### Problem

High background

### Potential Solution

This can either be due to non-specific staining due to antibodies cross-reaction or also PLA probes issues. Our experiments have validated mouse monoclonal antibody against Mfn1 (Abcam #ab57602), with the second antibody as a rabbit polyclonal antibody against Mfn2 (Abcam #ab50843). In myoblasts, antibodies were determined to work at a 1:200 dilution while in SkM antibodies were determined to work at a 1:50 dilution; however, specific conditions may affect the specific dilution needed. If all signal it too high, additional washing steps can aid in reducing probes. Alterations in reducing incubation time and blocking times may further reduce potential background. Beyond this, it is also important to include a control to ensure the probe sequences are highly specific to the target proteins and not causing additional background. Finally, if the background is more akin to coalescence, it is possible that amplification steps are too long, or overexposure is occurring.

### Problem

Low Signal

### Potential Solution

If possible, try to add a positive control, such as through overexpression of Mfn and Mfn2, which will likely cause increased mitochondria clustering and Mfn1-Mfn2 interactions ^15^. Traditional immunofluorescence can also serve as an effective positive control ^9^. Otherwise, attempt to increase concentrations of probes or antibodies, or while optimizing the washing steps and incubation time and temperature accordingly. Ensure all washes are done completely without any leftovers with WASH buffers at room temperature. It is especially important the ligation and amplification steps are at the correct temperature.

### Problem

Abundance of non-specific staining

### Potential Solution

One intriguing future avenue is the application of flag tags alongside PLA. Flag tags are short sequences that may be added to the N- or C-terminus and have a more unique peptide sequence that can aid in potential false positives ^13^. Coimmunoprecipitation can also be measured with Flag-tagged proteins by utilizing anti-FLAG magnetic beads ^16^. Thus, this can reduce the possibility of false positives and allow for a greater understanding of protein interactions, with coimmunoprecipitation also ensuring that protein interactions occur through other methods.

### Problem

Difficulty Applying Procedure

### Potential Solution

This may arise from troublesome sample preparation. Ensure all samples are properly fixed and the permeabilization solution is applied at the appropriate time.

## Resource Availability

### Lead contact

Further information and requests for resources and reagents should be directed to and will be fulfilled by the lead contact, Antentor Hinton.

### Materials availability

All generated materials, if applicable, are created in methods highlighted in the text above.

### Data and code availability

Full data utilized and requests for data and code availability should be directed to and will be fulfilled by the lead contact, Antentor Hinton.

## Financial & Competing Interests’ Disclosure

All authors have no competing interests.

This project was funded by the UNCF/Bristol-Myers Squibb E.E. Just Faculty Fund, BWF Career Awards at the Scientific Interface Award, BWF Ad-hoc Award, NIH Small Research Pilot Subaward to 5R25HL106365-12 from the National Institutes of Health PRIDE Program, DK020593, Vanderbilt Diabetes and Research Training Center for DRTC Alzheimer’s Disease Pilot & Feasibility Program. CZI Science Diversity Leadership grant number 2022-253529 from the Chan Zuckerberg Initiative DAF, an advised fund of Silicon Valley Community Foundation (to A.H.J.). NSF EES2112556, NSF EES1817282, NSF MCB1955975, and CZI Science Diversity Leadership grant number 2022-253614 from the Chan Zuckerberg Initiative DAF, an advised fund of Silicon Valley Community Foundation (to S.D.) and National Institutes of Health grant HD090061 and the Department of Veterans Affairs Office of Research award I01 BX005352 (to J.G.). Additional support was provided by the Vanderbilt Institute for Clinical and Translational Research program supported by the National Center for Research Resources, Grant UL1 RR024975–01, and the National Center for Advancing Translational Sciences, Grant 2 UL1 TR000445–06 and the Cell Imaging Shared Resource.

## Notes

### Competing Interest Statement

The authors have declared no competing interest.

### Summary of Updates

Updated figures and manuscript.

